# EssC is a specificity determinant for *Staphylococcus aureus* type VII secretion

**DOI:** 10.1101/252353

**Authors:** Franziska Jäger, Holger Kneuper, Tracy Palmer

## Abstract

The Type VII protein secretion system (T7SS) is found in actinobacteria and firmicutes, and plays important roles in virulence and interbacterial competition. A membrane-bound ATPase protein, EssC in *Staphylococcus aureus,* lies at the heart of the secretion machinery. The EssC protein from *S. aureus* strains can be grouped into four variants (EssC1-EssC4) that display sequence variability in the C-terminal region. Here we show that the EssC2, EssC3 and EssC4 variants can be produced in a strain deleted for *essC1* and that they are able to mediate secretion of EsxA, an essential component of the secretion apparatus. They are, however, unable to support secretion of the substrate protein EsxC, which is encoded only in essC1-specific strains. This finding indicates that EssC is a specificity determinant for T7 protein secretion. Our results support a model where the C-terminal domain of EssC interacts with substrate proteins whereas EsxA interacts elsewhere.

---

The type VII secretion system (T7SS) is found primarily in bacteria of the actinobacteria and firmicutes phyla and secretes proteins that lack cleavable N-terminal signal peptides. The system is best characterised in mycobacteria, where it is designated ESX, and pathogenic members of the genus can encode up to five copies of the secretion machinery (1, 2). Substrates of the T7SS may vary in size but are usually α-helical in nature. Every T7SS analysed to date secretes at least one protein of the WXG100 superfamily. Proteins of this family are small helical hairpins that have a conserved W-X-G amino acid motif in a short loop between the two helices (3, 4). A YxxxD/E motif, located at the C-termini of some WXG100 proteins acts, in concert with the WXG motif, as a bi-partite targeting sequence for T7 secretion (5-8). WXG100 proteins are secreted as folded dimers; in actinobacteria these are heterodimers of paired WXG100 proteins whereas in firmicutes these may also be homodimers (8). The T7SS also secretes much larger substrates that share a similar four-helical bundle arrangement of the WXG100 protein dimers (7, 9, 10). Some T7 substrates interact with chaperones prior to secretion and there is evidence that secretion of LXG domain substrates in firmicutes is dependent on complex formation with a WXG100 protein partner (11-13).

There are commonalities and differences between the T7SS of actinobacteria and firmicutes (14). A membrane-embedded ATPase of the FtsK/SpoIIIE family termed EccC/EssC is found in all T7SSs. In both systems the protein shares a similar overall topology, with two transmembrane domains that are usually followed by three P-loop ATPase domains at the C-terminus. Although all three P-loop ATPase domains are capable of binding ATP, mutagenesis studies have indicated that only ATP hydrolysis by domain 1 is essential for T7 secretion (15, 16). In actinobacteria, a hexameric arrangement of the EccC ATPase lies at the centre of a 1.8MDa complex that also contains six copies of the EccB, EssD and EccE proteins (17). In firmicutes, homologues of EccB, D and E are absent and a distinct set of membrane proteins, EsaA, EssA and EssB, work alongside the ATPase, EssC, to mediate T7 secretion (18-22). In *Staphylococcus aureus* and *Bacillus subtilis* a secreted WXG protein, EsxA, and a small cytoplasmic protein, EsaB, are also required for T7SS activity (18,19, 21-23) (Fig 1A).

**Figure 1.**
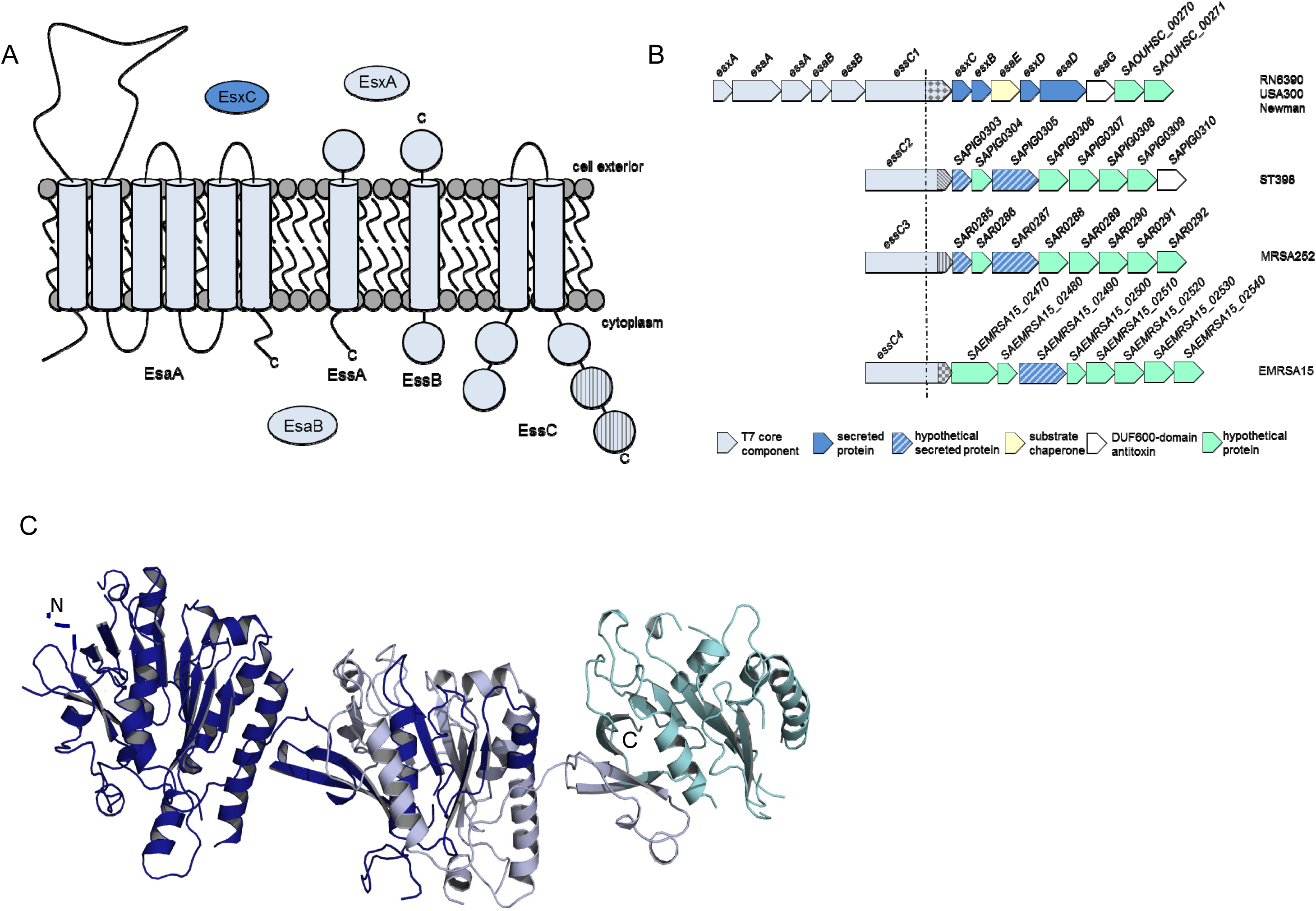
Sequence variability in *S. aureus* EssC. A. The *S. aureus* T7 secretion machinery. Components that are essential for T7 secretion are shown in light grey with their subcellular locations. The hatched domains of EssC indicate sequence-variable regions. The substrate protein EsxC, found only in strains with the EssC1 variant, is shown in blue. B. Genetic organisation of the *S. aureus ess* locus in the four different ess strain variants. Since the 3’ boundaries of the ess loci are not known, the first eight genes downstream of *essC* are shown in each case. The dotted line indicates the approximate position of *essC* sequence divergence and the shading at the 3’ end of *essC* represents the region of sequence variability. C. Structural model of the ATPase domains of *S. aureus* EssC (generated using amino acids 601-1078 of EMRSA15 EssC) using Phyre2 (www.sbg.bio.ic.ac.uk/∼phyre) with the structure of EccC from *Thermomonospora curvata* (16) as a template. The shading is dark blue: residues 601-1078, very highly conserved; light blue: residues 1079-1289 (where the EssC1 sequence diverges from the remaining EssC); cyan: residues 1290-1479 (variable C-terminal region).

The EccC/EssC ATPase has previously been implicated in substrate recognition. Crosslinking and co-purification experiments have identified complexes of *S. aureus* EssC with substrates EsaD (also called EssD) and EsxC (12, 24), and the EccC ATPase domains have been co-crystallised with a peptide from the C-terminus of the WXG protein, EsxB (16). Further evidence in support of a role for EssC in substrate recognition comes from genomic analysis of *S. aureus* (25). It was noted that there was sequence variability at the ess locus across different *S. aureus* strains. Genes coding for the core components EsxA-EssB are highly conserved (Fig 1B), as is the 5’ end of *essC,* but the 3’ portion of the gene falls into one of four sequence groupings (25). The *essC* sequence type strictly co-varies with the sequence of adjacent 3’ genes, some of which are known or strongly predicted to encode secreted substrates. This would be consistent with the C-terminal variable region of EssC playing a role in substrate recognition. In this study we have addressed this hypothesis directly by assessing whether EssC proteins from the EssC2, EssC3 and EssC4 classes can support the secretion of the EssC1 substrate, EsxC (26) and of the core component, EsxA.

*S. aureus* EssC proteins are approximately 1480 amino acids in length and have a common domain organisation, with two forkhead associated (FHA) domains at their N-termini, two transmembrane domains and three repeats of a P-loop ATPase domain at their C-termini (27, 28; Fig 1A). Sequence analysis indicates that *S. aureus* EssC proteins are almost sequence invariant until part way through the second ATPase domain, where the EssC1 variant, found in strains such as RN6390, Newman and USA300 starts to diverge (Fig 1C; Fig S1). The EssC2, EssC3 and EssC4 variants are more similar to one another, and share almost identical sequence until ATPase domain 3 where they also start to vary (Fig 1C; Fig S1). Of the four ATPases, variants 2 (from strain ST398) and 3 (from strain MRSA252) are the most similar (Fig S1).

We have previously constructed an in-frame deletion of *essC* in strain RN6390 and shown that this results in the inability to export both the core machinery component, EsxA, and the substrates EsxC and EsaD (12, 19). This secretion deficiency could be rectified by re-introduction of EssC1 encoded on plasmid pRMC2 (29). Fig 2A shows that production of EssC1 could be also restored when it was encoded on the expression vector pRAB11 (30), and that re-introduction of plasmid-encoded EssC1 resulted in strong secretion of both EsxA and EsxC in the RN6390 Δ*essC* strain.

**Figure 2.**
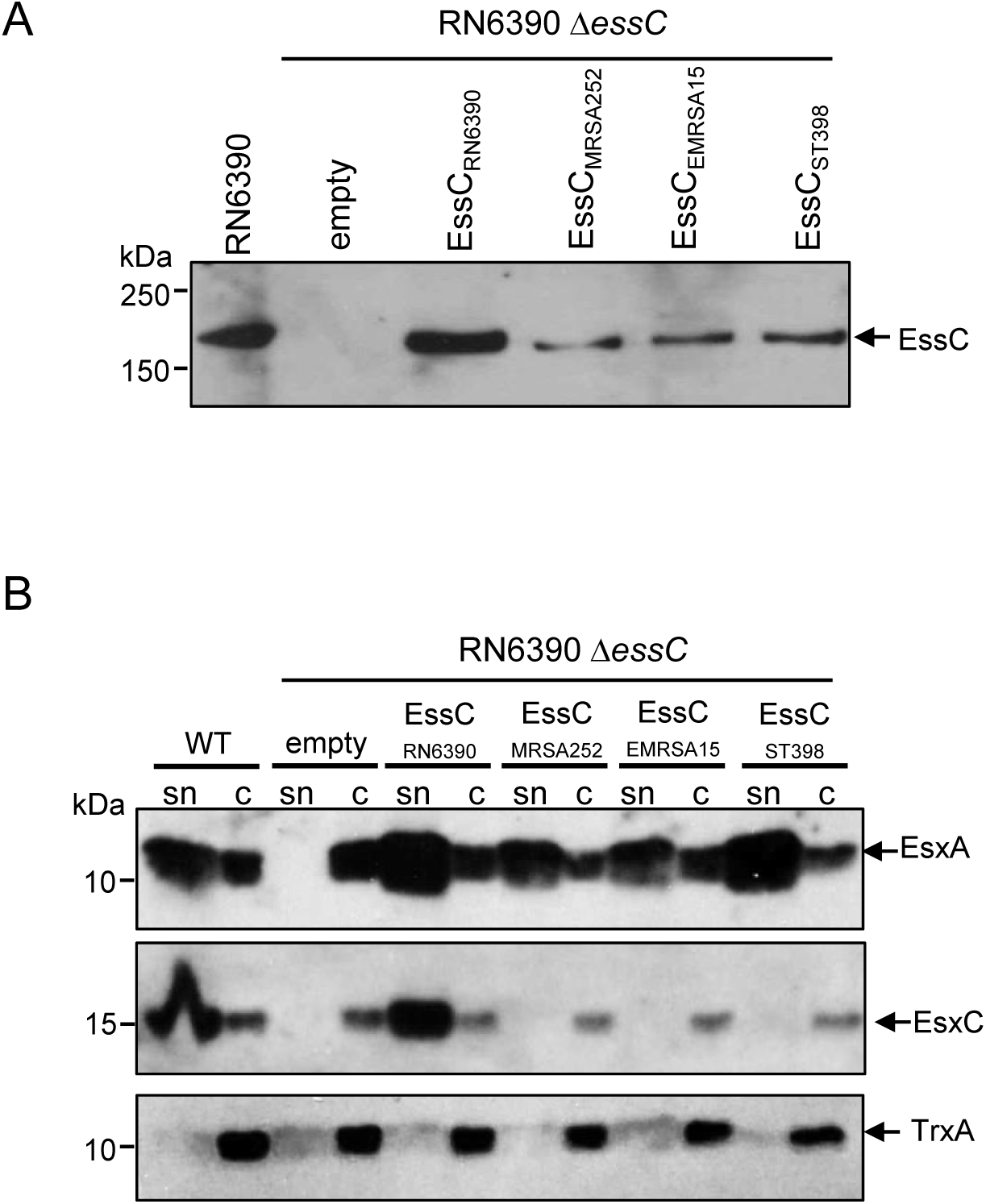
Non-cognate EssC variants support secretion of EsxA but not EsxC. A and B. Strain RN6390 or the isogenic *essC* deletion strain carrying pRAB11 (empty) or pRAB11 encoding the indicated *essC* variant was subcultured into TSB medium supplemented with 1 nM hemin (32) and either 25ng/ml (RN6390 Δ*essC*/pEssC_RN6390_) or 100ng/ml (RN6390 Δ*essC*/pEssC_MRSA252_/pEssC_ST398_/pEssC_EMRSA15_) anhydrotetracycline (ATC), to induce plasmid-encoded gene expression. Strain were grown aerobically until an OD_600_ of 2 was reached after which A. 10 μl of OD_600_ 1 adjusted cells were separated on an 8% bis-Tris acrylamide gel and analysed by western blotting using anti-*essC* antisera (19), or B. cultures were separated into supernatant and whole cell fractions and equivalent of 200 μl of culture supernatant (sn) and 10 μl of resuspended cell sample (c) adjusted to an OD_6_00 = 1 were separated on a 15 % bis-Tris-gel and immunoblotted using the antiserum raised against EsxA (19), EsxC (19) or the cytosolic control TrxA (33).

Next, we amplified the genes for **essC*2* (from strain ST398), **essC*3* (from strain MRSA252) and *ess4* (from strain EMRSA15) and also cloned these into pRAB11 (see Table S1 for oligonucleotides used for these experiments). We first confirmed that the three variant *essC* proteins could be stably produced in the RN6390 Δ*essC* strain background. To this end anhydrotetracycline (ATC) was added to induce plasmid-encoded production of *essC* and whole cell samples were analysed by blotting with an *essC* antiserum. It should be noted that the antiserum used was raised against a truncated protein covering the last two ATPase domains of the *essC*1 variant (19). As shown in Fig 2A, each of the *essC*2, *essC*3 and *essC*4 variants could be recognised by this antibody, but not so strongly as the cognate *essC*1 due to a lack of conservation of epitopes in this region of the protein. We conclude that all *essC* variants can be produced in strain RN6390.

Next, we asked whether the variant *essC* proteins in RN6390 could support T7 protein secretion. Fig 2B (top panel) shows that secretion of the EsxA core component was indeed supported by each of these *essC* proteins, indicating that each *essC* variant was functional in the heterologous strain background. However, none of the *essC* variants were able to support secretion of the substrate protein, EsxC. Taken together these results confirm that *essC* is a specificity determinant for substrate secretion by the *S. aureus* T7SS. The findings strongly suggest that the sequence invariant regions of *essC* proteins are involved in mediating interactions with the conserved T7 core components, including the secreted protein EsxA (which has >99% sequence identity across all sequenced *S. aureus* strains) and that the sequence variable region, primarily ATPase domain 3, is involved in substrate recognition. This might imply that EsxA and EsxC are secreted by different mechanisms.

Finally, it is interesting to note that secretion of all known substrates mediated by the *essC*1 variant is dependent on a chaperone protein, EsaE/EssE (12, 24). Some substrates of the actinobacterial T7SS also interact with specific chaperones of the EspG family to ensure delivery to the cognate secretion machinery (11, 31), although other substrates appear to be exported independently of a specific chaperone (2). No protein with any detectable sequence homology to either EsaE or EspG is encoded at the ess loci of the **essC*2, *essC*3* or **essC*4* strain variants. In future it will be interesting to determine whether the mechanism of substrate targeting differs across the Ess subtypes in *S. aureus.*

## ACKNOWLEDGEMENTS

Dr Jon Cherry is thanked for his help with generating the structural model in Fig 1C. This study was supported by the Wellcome Trust (through Investigator Award 10183/Z/15/Z to TP, the Biotechnology and Biological Sciences Research Council (through the EASTBIO Doctoral Training Partnership award number BB/J01446X/1 which provided a PhD studentship (to FJ), and through grant BB/H007571/1) and by Medical Research Council (through grant MR/M011224/1). The authors declare no conflicts of interest.

---

**Table S1.**
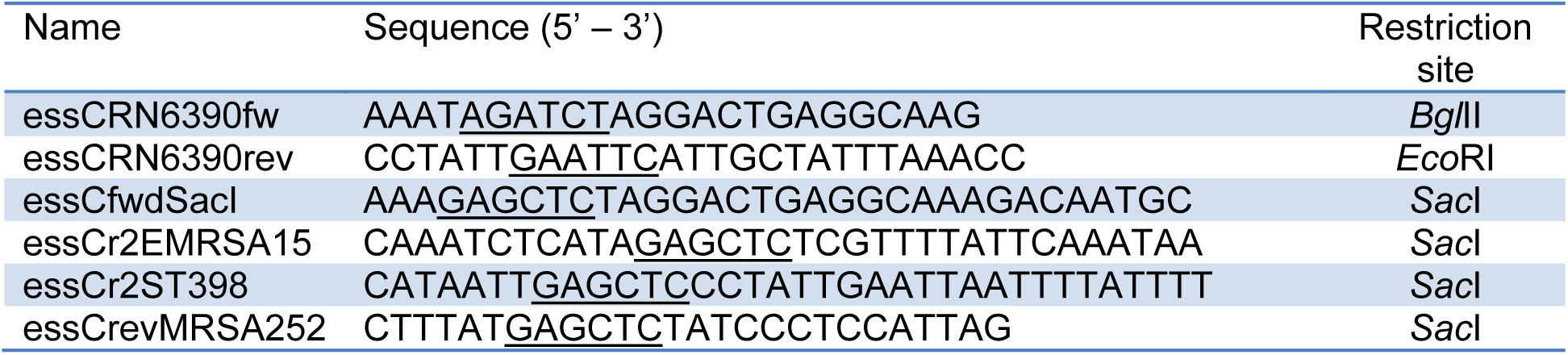
Oligonucleotides used in this study. The **essC** gene from RN6390 was amplified using oligonucleotides *essC*RN6390fw and *essC*RN6390rev and cloned into plasmid pRAB11 as a *BglII-EcoRI* fragment. The other three *essC* genes were amplified using the same forward primer (*essC*fwdSacI) and a strain-specific reverse primer and cloned into pRAB11 as SacI fragments. All inserts were confirmed for directionality and fidelity by DNA sequencing.

**Figure S1.**
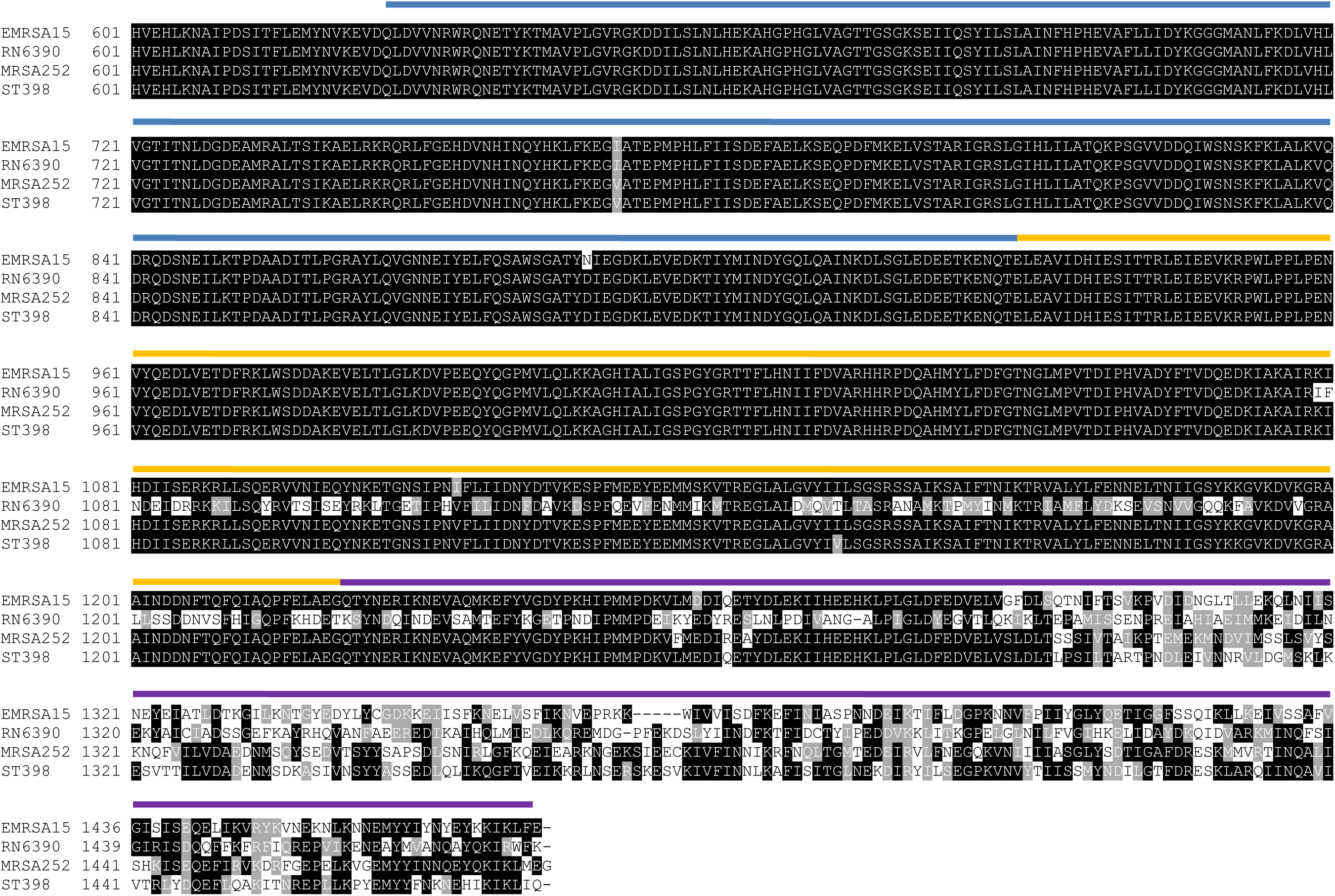
Alignment of *essC* sequences from the indicated *S. aureus* strains. The alignment was generated using Clustal W (http://www.ch.embnet.org/software/ClustalW.html) and shaded using Boxshade (https://embnet.vital-it.ch/software/BOX_form.html) and is shown from amino acid 600 onwards. The blue, yellow and purple lines above the alignment delimits the extend of ATPase domains 1,2 and 3, respectively.

